# Developmental Stage and Cellular Context Determine Oncogenic and Molecular Outcomes of *Ezh2*^Y641F^ Mutation in Hematopoiesis

**DOI:** 10.1101/2024.11.14.622807

**Authors:** Sarah M. Zimmerman, Samantha J. Procasky, Sofia R. Smith, Jie-Yu Liu, Chad Torrice, George P. Souroullas

## Abstract

Mutations in the histone methyltransferase EZH2, particularly the Y641 hotspot mutation, have been implicated in hematologic malignancies, yet the effect of timing and cellular context on their oncogenic potential has remained unknown. In this study, we utilized a conditional allele with tissue-specific Cre drivers to investigate the effects of *Ezh2*^Y641F^ mutations at various stages of development, with a focus on the hematopoietic system. We found that ubiquitous heterozygous *Ezh2*^Y641F^ expression at birth, or conditional expression in hematopoietic or mesenchymal stem cells, led to decreased survival due to hematopoietic defects and bone marrow failure, with no evidence of malignancy. In contrast, *Ezh2*^Y641F^ expression in committed B cells drives lymphoma formation, highlighting the lineage-specific oncogenic activity of the mutation. Transcriptomic analysis of B cell progenitors revealed key pathway alterations between Cre models such as altered IL2-Stat5 signaling pathway, differential expression of E2F targets, and altered GTPase pathway expression driven by upregulation of Guanylate Binding Proteins (GBPs) in Mx1-Cre *Ezh2*^Y641F^ pro-B cells. We further found that the GBP locus is regulated by Ezh2-mediated H3K27me3, it is associated with poorer survival in Acute Myeloid Leukemia patients and has variable effects on apoptosis in human lymphoma and leukemia cell lines. These findings suggest that the *Ezh2*^Y641F^ mutation may alter immune regulatory pathways, cell differentiation and apoptosis, with potential implications for disease progression. Our results highlight the critical role of mutation timing and cellular context in EZH2-driven hematopoietic disease, resulting in distinct downstream changes that shape the oncogenic impact of EZH2.

## Introduction

Mutations in the EZH2 gene, a key regulator of chromatin structure and gene expression, play crucial roles in both normal development and malignancy. EZH2 is the enzymatic subunit of the Polycomb Repressive Complex 2 (PRC2) and mediates the methylation of lysine 27 on histone 3 (H3K27), contributing to chromatin compaction and transcriptional repression. The role of EZH2 in cancer is context-dependent, acting as either a tumor suppressor or oncogene depending on the cellular environment. For example, *EZH2* deletions are often observed in T cell acute lymphoblastic leukemia or acute myeloid leukemia, where it typically functions as a tumor suppressor ^1–5^. In contrast, overexpression or amplifications of EZH2 are oncogenic in many solid tumors, including breast, lung, prostate and melanoma ^6–10^.

A recurrent point mutation, *EZH2*^Y641F^ (or Y646, depending on isoform), in the SET catalytic domain is observed in 15–20% of mature B-cell lymphomas such as follicular lymphoma (FL) and diffuse large B-cell lymphoma (DLBCL) ^11,12^. Mouse models of the *Ezh2*^Y641F^ mutation have shown that Ezh2 is required for Germinal Center (GC) formation, and that expression of the mutant *Ezh2*^Y641F^ promotes GC hyperplasia and the development of B cell lymphoma ^6,8,13–15^. At the molecular level, *Ezh2*^Y641^ mutations were initially thought to be loss-of-function mutations ^11^, however, subsequent studies showed that they exhibit neomorphic activity, preferentially methylating mono- or di-methylated H3K27, but requiring the presence of wild-type EZH2 for initial mono-methylation steps ^16,17^. These observations are consistent with the fact that all *EZH2*^Y641^ mutations identified to date in human patients are heterozygous^12^. Globally, these mutations alter the distribution of H3K27me3, with increased deposition in some regions, decreased in others, and spreading of H3K27me3 past canonical boundaries ^6,14,18^. These findings suggest that *EZH2*^Y641F^ mutations are not purely gain-of-function but likely neomorphic, and distinct from simple overexpression of wild-type *EZH2*.

While the oncogenic properties of *Ezh2*^Y641^ mutations have been well studied in mature B cell lymphoma, they also occur in other malignancies, including B cell lymphoblastic leukemias, acute myeloid leukemia (AML), and solid tumors such as melanoma, bone, breast, mesothelioma and endometrial cancers (**Fig. 1a, b**). While Ezh2 also plays a critical role in hematopoietic stem cell functions ^19–21^, it is surprising that these mutations are infrequent in non-B cell malignancies (**Fig. 1a, b**), implying that the cell of origin and developmental timing significantly influence their oncogenic potential.

**Figure 1.**
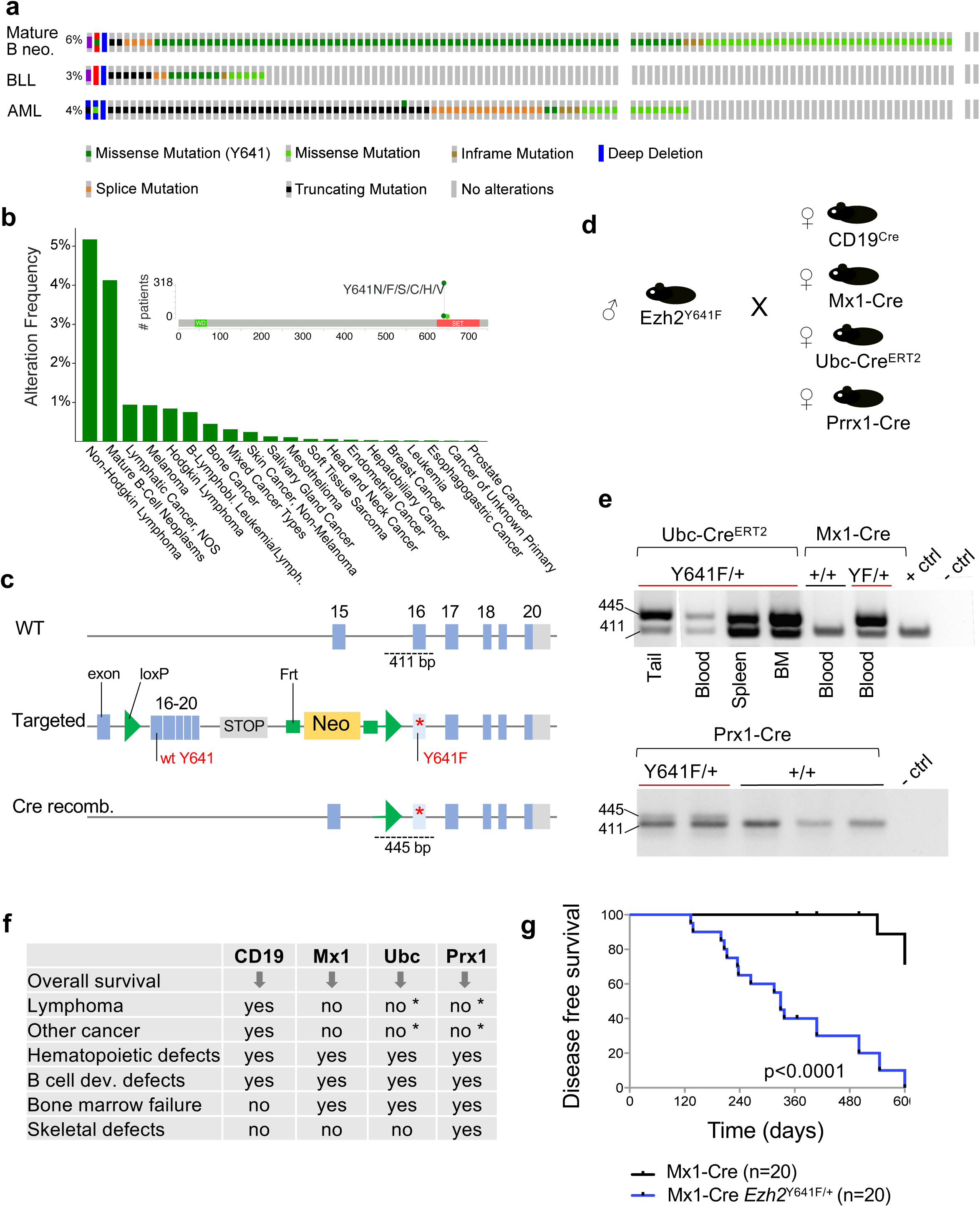
Expression of *Ezh2*^Y641F^ in hematopoietic stem cells shortens overall survival. **a.** Genetic alterations in EZH2 across mature B cell neoplasms, B cell leukemias (BLL), and acute myeloid leukemia (AML), analyzed using TCGA data from the Genie cohort (v16.1) on cBioPortal. B cell neoplasms, n=6444; BLL, n=911; AML, n=3859. **b.** Frequency of *EZH2*^Y641F^ mutations across various cancer types, based on Genie cohort (v16.1) data analyzed via cBioPortal. **c.** Schematic of Cre-lox strategy used to induce *Ezh2*^Y641F^ expression in mice. **d.** Breeding strategy for cell type-specific expression of *Ezh2*^Y641F^ using different Cre drivers. **e.** PCR confirmation of Ezh2 recombination in specific tissues by each Cre allele. **f.** Summary of phenotypes observed in each Cre-driven *Ezh2*^Y641F^ expression model. **g.** Disease-free survival curves for Mx1-Cre *Ezh2*^Y641F^ compared to Cre-only controls (log-rank test p<0.0001); median survival for Mx1-Cre *Ezh2*^Y641F^ mice is 329 days.

Mechanistically, *Ezh2*^Y641^ mutations have been associated with diverse pathways. In lymphoma models, *Ezh2*^Y641F^ mutations are linked to disrupted differentiation programs ^8^, Myc transcriptional signatures ^6,22^, interference with p53-mediated mTORC1 signaling ^23^, and cooperation with Bcl2 and Bcl6 ^6,8,13^. In melanoma, *Ezh2*^Y641F^ expression accelerates tumorigenesis in cooperation with oncogenic *Braf*^V600E^ mutations, while also modulating anti-tumor immunity through interferon signaling ^6,24–26^. These findings underscore the context-dependent effects of *EZH2*^Y641F^, which vary significantly based on cell type and developmental stage.

Somatic mutations in EZH2, including those in the SET domain, have also been identified in patients with Weaver syndrome, a rare overgrowth disorder characterized by accelerated skeletal maturation, intellectual disability and increased cancer susceptibility ^27–30^. While most mutations so far characterized in patients tend to be missense or truncating mutations, mouse models suggest that EZH2 activity modulates growth phenotypes through its effect on H3K27me3.

To address the gaps in understanding how developmental stage and cell of origin influence the phenotypic and oncogenic outcomes of *Ezh2*^Y641F^ mutations, we investigate the effects of this mutation in a range of contexts using tissue- and cell type-specific Cre driver alleles. By focusing on phenotypic and molecular differences across stages of hematopoietic development and tumorigenesis, this study explores how the timing and context of *EZH2*^Y641F^ mutations shape their downstream effects, offering novel insights into their role in hematopoiesis and cancer.

## Materials and Methods

### Mice

All mice were backcrossed to the C57Bl/6 background and housed in an Association for Assessment and Accreditation of Laboratory Animal Care (AAALAC)-accredited facility and treated in accordance with protocols approved by the Institutional Animal Care and Use Committee (IACUC) for animal research initially at the University of North Carolina-Chapel Hill, and later at Washington University in St. Louis. Both male and female mice were included in all experiments.

### Genotyping and induction of *Ezh2*^Y641F^ allele recombination

Genomic DNA was extracted from mouse tails and genotyped for Cre and *Ezh2*^Y641F^ alleles using standard PCR protocols. Induction of the Mx-Cre allele was achieved by intraperitoneal injection of polyinosinic-polycytidylic acid (polyI:C) (Sigma P0913), three injections every other day at 300 μg per dose. Induction of the Ubc-Cre^ERT2^ allele was induced by intraperitoneal injections of tamoxifen dissolved in corn oil at approximately 75 mg tamoxifen/kg body weight, at 2- and 4- days post birth. Tissue-specific CD19- and Prx1-Cre alleles are not inducible. Recombination of the *Ezh2*^Y641F^ allele was confirmed by PCR. Primer sequences are provided in Supplementary Table 2.

### Competitive bone marrow transplantation assays

Bone marrow was collected from the femora and tibiae for donor mice. Transplant recipient mice were lethally irradiated with a split dose of 10.5 Gy, 5 hours apart, and injected intravenously with a mixture of C57Bl/6 CD45.2 transgenic donor whole bone marrow and CD45.1 wild-type competitor bone marrow, 250K cells each. Both male and female mice were used as transplant recipients, all 8 weeks old.

### Flow cytometry

Peripheral blood was collected into tubes containing potassium EDTA and complete blood counts were counted on a Hemavet (Drew Scientific). Flow cytometry analysis was carried out on LSRII (BD), Dako Cyan (Beckman Coulter) and Attune (Thermo) using antibodies listed in Supp. Table 1. Data was analyzed using FlowJo software (TreeStar Inc.).

### Histology

Tissue samples were fixed in 10% formalin, processed for paraffin embedding by Histowiz, sectioned at 5 μm and stained with hematoxylin and eosin (H&E). Bones were decalcified using 10% EDTA pH7.2 for two weeks at 4°C and processed similarly for bone analysis.

### RNA sequencing and analysis

Pro- and pre-pro-B cells were sorted from bone marrow using fluorescence-activated cell sorting (FACS) and processed for RNA extraction using Qiagen RNeasy Micro kit (Qiagen 74004). Libraries were prepared and sequenced on an Illumina NovaSeq 6000 platform. Differential gene expression analysis was performed using EdgeR and Limma, and pathway analysis utilized GAGE and WGCNA. RNA-seq data are available in GEO (accession number GSE270480).

### Cell culture and generation of GBP2 overexpression cell lines

Cells were grown in RPMI medium 1640 (Gibco Cat# 11875-093) with 10% FBS (Millipore Sigma Cat# F0926) and 1% penicillin-streptomycin (Genesee Scientific Cat# 25-512). The GBP2 lentiviral vector (Vector Builder ID VB900000-0734xke) was used to overexpress human GBP2, and the control vector is TLCV2 (Addgene 87360) with no sgRNA inserted. Lentivirus was produced in 293T cells via transfection with PEI.

### Statistical Analysis

Statistical analysis was performed using GraphPad Prism. Bar graphs represent mean and either standard deviation or standard error of the mean as indicated in the figure legends. Sample sizes were determined based on pilot studies or historical data from similar experiments. Power analysis was performed for two-tailed analysis, under the assumption of normal distribution, with a significance level of 0.05 and power of 0.8. Differences between groups were determined using Student’s t-test unless otherwise indicated in the figure legend.

## Results

### Heterozygous expression of *Ezh2*^Y641F^ during early embryonic development results in developmental and hematopoietic defects and decreased survival

To investigate the oncogenic potential of *Ezh2*^Y641F^, we used a conditional *Ezh2*^Y641F^ allele ^6^ activated by tissue-specific Cre drivers (**Fig. 1c**). We crossed the *Ezh2*^Y641F^ allele to four Cre drivers: ubiquitously expressed and tamoxifen-inducible Ubc-Cre^ERT2^ ^31^, limb-bud-specific Prx1-Cre ^32^, hematopoietic-specific Mx1-Cre^33^, and B cell-specific CD19Cre **(Fig. 1d)**. Since germline deletion of Ezh2 in the mouse is lethal, and *Ezh2*^Y641F^ mutations are always heterozygous in patients, we maintained these animals in a heterozygous state and confirmed proper allele recombination in multiple tissues (**Fig. 1e**).

Across all models, *Ezh2*^Y641F^ expression resulted in reduced survival, with bone marrow failure observed in all crosses except CD19-Cre, which uniquely developed B-cell, (**Supp. Fig. 1a**) consistent with our previous findings ^6^. The Prx1-Cre crosses exhibited lower body weight and joint-related skeletal defects (**Supp. Fig. 1b**) as evident by abnormal animal posture and locomotion. Mice in the Ubc-Cre^ERT2^, Prx1-Cre and Mx1-Cre groups exhibited very similar hematopoietic defects such as anemia, enlarged spleens and reduced peripheral B cell output (**Fig. 1f**, **Supp. Fig. 1b-f**). Interestingly, none of these crosses resulted in the development of B cell lymphoma, or other malignancies, but instead showed signs of bone marrow failure by six months of age. Bone marrow failure and decreased survival limited further observation of other potential malignancies requiring longer latency periods.

Given these outcomes, we focused our detailed investigation on the two hematopoietic-specific Cre drivers and hypothesized that the cell of origin during hematopoiesis critically determines the oncogenic potential of the *Ezh2*^Y641F^ mutation.

### Expression of *Ezh2*^Y641F^ in HSCs impairs B cell development and peripheral output

Given the absence of overt malignancy in the Mx1-Cre *Ezh2*^Y641F^, we analyzed hematopoiesis across multiple organs over time. In young mice (4-10 weeks old), flow cytometric analysis of peripheral blood revealed no changes in lymphoid or myeloid populations. However, as the mice aged, they exhibited increased mortality compared to Mx1-Cre *Ezh2*^WT^ control animals with a median survival of 329 days (**Fig. 1g**). Additionally, older animals exhibited progressive anemia, thrombocytopenia and lymphopenia (**Fig. 2a-b)**. Flow cytometry revealed a significant reduction in peripheral B cells, while other lymphoid populations were largely unaffected (**Fig. 2c-e**). To determine if defects in B cell differentiation in the bone marrow contributed to this phenotype, we analyzed B cell progenitor populations in the bone marrow of young animals. We identified a partial block in B cell development at the pre-pro to pro B cell transition (**Fig. 2f-g**). These findings indicate that expression of mutant *Ezh2*^Y641F^ in hematopoietic stem and progenitor cells disrupts B cell development, resulting in reduced mature B cell output in the peripheral blood.

**Figure 2.**
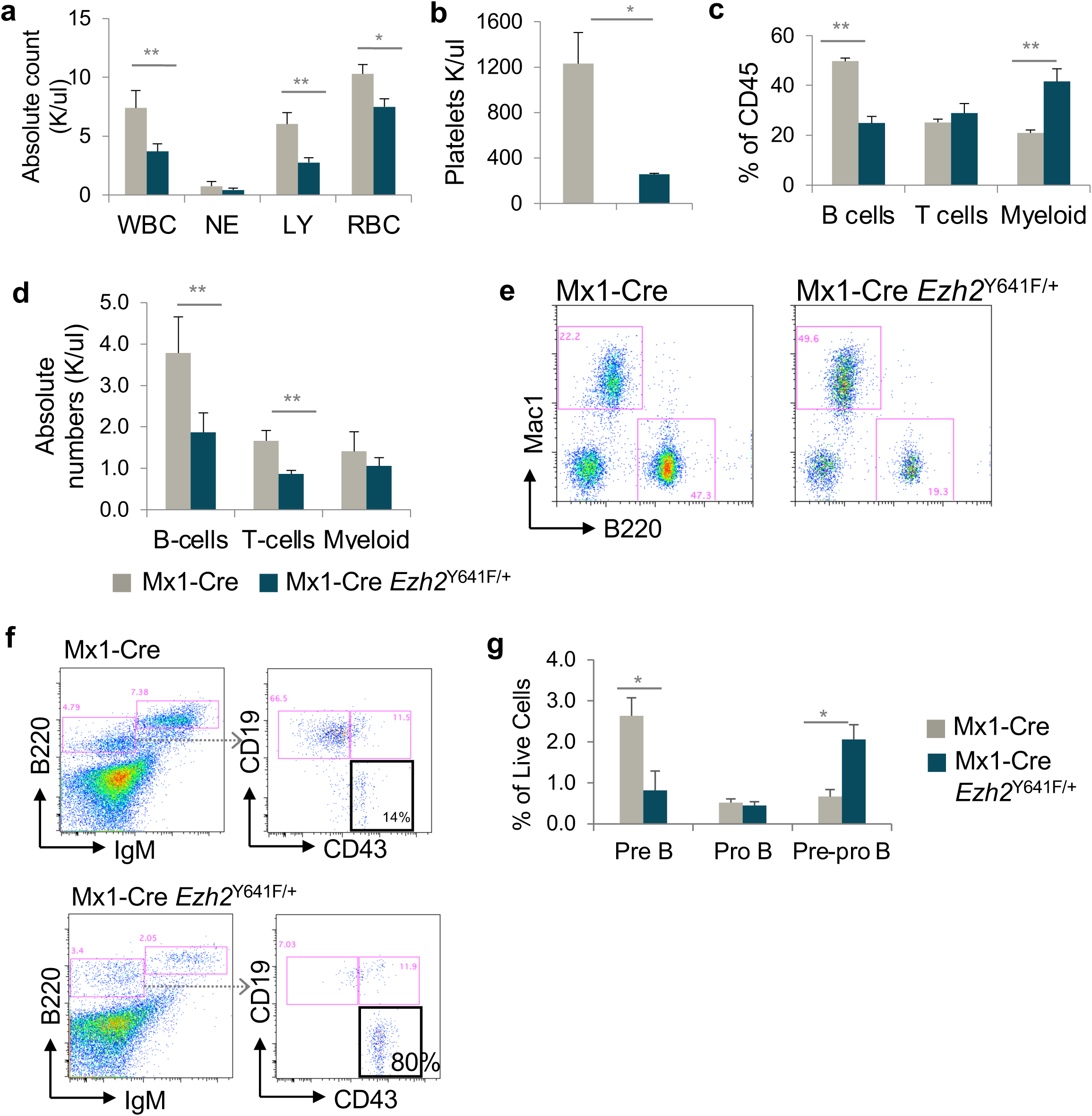
Partial block in B cell development in Mx1-Cre *Ezh2*^Y641F^ mice. **a.** Complete peripheral blood cell counts (CBC) showing white blood cells (WBC), neutrophils (NE), lymphocytes (LY), red blood cells (RBC), and platelets in peripheral blood from Mx1-Cre *Ezh2*^Y641F^ and Mx1-Cre control mice, n>5 per group **b.** Platelet counts from CBC analysis. **c.** Flow cytometry analysis of peripheral blood in Mx1-Cre *Ezh2*^Y641F^ and Mx1-Cre only control mice, n>5. **d.** Calculated absolute numbers of B cells, T cells and Myeloid cells in the peripheral of Mx1-Cre *Ezh2*^Y641F^ and Mx1-Cre control from flow cytometry (panel a) and CBC (panel c). **e.** Representative flow plots of data in c. **f.** Representative flow cytometry of bone marrow B cell progenitor populations in Mx1-Cre *Ezh2*^Y641F^ and Mx1-Cre only control mice; black box indicates pre-pro B cells. **g.** Quantification of bone marrow B cell populations in Mx1-Cre *Ezh2*^Y641F^ and Mx1-Cre control mice, n>5. Error bars indicate standard deviation. * p<0.01, **p<0.001.

### Expression of *Ezh2*^Y641F^ in HSCs results in skewed hematopoietic progenitor populations, high bone mass and bone marrow failure

Given the oncogenic role of *Ezh2*^Y641F^ mutations in lymphoma, we hypothesized that the observed phenotypes in Mx1-Cre *Ezh2^Y641F^*mice might be driven by an underlying hematopoietic malignancy. To explore this hypothesis, we first analyzed hematopoietic progenitor populations in the bone marrow. In young mice (2-3 months old), we did not find any significant differences between Mx1-Cre *Ezh2^WT^* and Mx1-Cre *Ezh2*^Y641F^ (data not shown). However, by six months of age, mice expressing *Ezh2*^Y641F^ exhibited decreased bone marrow cellularity (**Fig. 3a**), accompanied by significant skewing of hematopoietic progenitors towards proportionally more common lymphoid progenitors and less myeloid progenitors (**Fig. 3b-d**). Notably, the bones of *Ezh2*^Y641F^ mice were pale, consistent with anemia and decreased hematopoietic production (**Fig. 3e**). Histological analysis did not find any evidence of blasts or hematopoietic malignancy, but instead revealed increased trabecular bone mass and decreased bone marrow cellularity, suggesting altered bone remodeling (**Fig. 3f**). Although Cre expression under the Mx-1 promoter is mostly restricted to hematopoietic populations, its expression has been detected in other tissues, such as osteoclasts, which are derived from myeloid precursors ^35^. This raises the possibility that *Ezh2*^Y641F^ expression in osteoclasts may disrupt bone resorption, leading to increased bone mass. This altered bone structure could subsequently constrain the bone marrow microenvironment, contributing to bone marrow failure.

**Figure 3.**
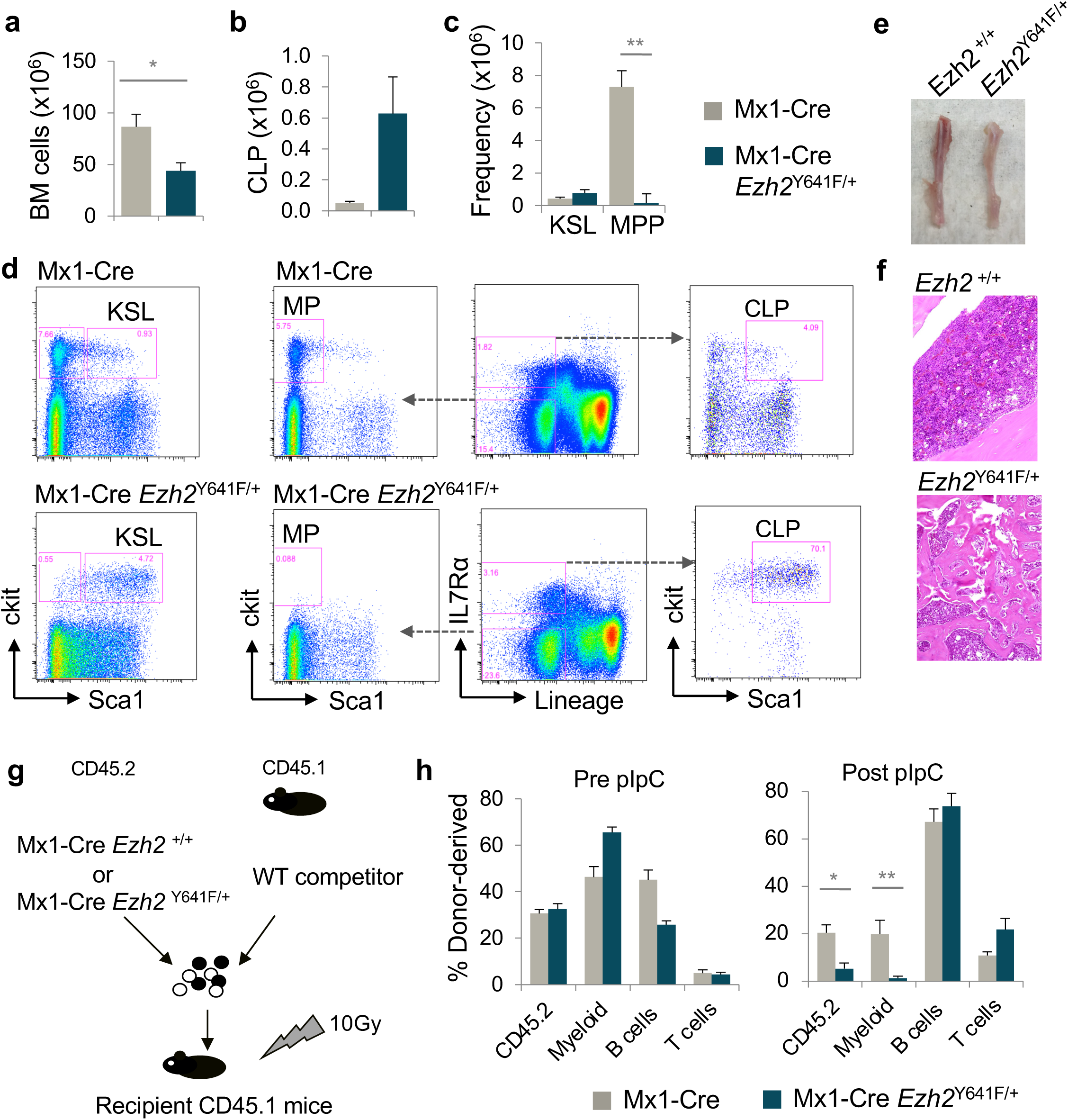
Defective cell intrinsic hematopoietic stem cell function after induced expression of *Ezh2*^Y641F^. **a.** Total bone marrow cellularity in Mx1-Cre *Ezh2*^Y641F^ mice and Mx1-Cre control mice, n>5 per group. **b.** Common Lymphoid Progenitors (CLP) counts in bone marrow of Mx1-Cre *Ezh2*^Y641F^ and Mx1-Cre only control mice, n>5. **c.** Analysis of ckit+ Sca1+ Lineage negative (KSL) and multipotent progenitor (MP) populations. **d.** Representative flow cytometry plots of bone marrow hematopoietic progenitor populations. **e.** Image of tibiae from a Mx1-Cre *Ezh2*^Y641F^ mouse (right) and control (left). **f.** H&E-stained bone sections from Mx1-Cre *Ezh2*^Y641F^ (bottom) and Mx1-Cre control mice (top). **g.** Workflow for competitive bone marrow transplant assay with the indicated genotypes. **h.** Donor cell engraftment analysis post competitive bone marrow transplant from, n=8 mice per group. Error bars represent standard deviation, * p<0.01, **p<0.001.

### Expression of *Ezh2*^Y641F^ in HSCs disrupts cell intrinsic stem cell functions

The absence of malignant transformation in Mx1-Cre *Ezh2*^Y641F^ animals was unexpected, particularly given that *Ezh2*^Y641F^ expression persisted across all B cells, and CD19-Cre *Ezh2*^Y641F^ mice developed B cell neoplasia before the onset of bone marrow failure in Mx1-Cre mice.^6^ One possible explanation is that early changes to the bone marrow microenvironment restrict B cell development and output, thus reducing the likelihood of acquiring secondary mutations necessary for B cell transformation.

To investigate this possibility, we performed bone marrow transplantation assays using hematopoietic stem and progenitor cells isolated from Mx1-Cre *Ezh2*^Y641F^ and Mx1-Cre control mice. These assays were performed using donor bone marrow from young animals (∼8 weeks old), prior to any detectable defects in progenitors or cellularity as described above. Equal numbers of test donor (CD45.2) and competitor (CD45.1) bone marrow cells (500K) were transplanted into lethally irradiated recipient mice (CD45.1) (**Fig. 3g**). Four weeks after the transplants, we injected mice with 13 mg/kg of poly(I:C) every other day and analyzed peripheral blood chimerism starting at two weeks after the last dose. Peripheral blood analysis by flow cytometry before Cre induction with poly(I:C) showed no significant differences in engraftment or differentiation. However, by 24 weeks post Cre induction, recipient mice receiving *Mx1-Cre Ezh2*^Y641F^ HSCs exhibited a marked decline in peripheral blood chimerism and impaired myeloid differentiation (**Fig. 3h**), indicating a loss of self-renewal and differentiation capacity. Long term monitoring of these animals did not reveal any evidence of disease.

### Extramedullary hematopoiesis in the spleens of Mx1-Cre *Ezh2*^Y641F^ mice

One of the most prominent phenotypes of Mx1-Cre *Ezh2*^Y641F^ mice was splenomegaly (**Fig. 4a-b**). To determine whether this phenotype was associated with a splenic malignancy, we performed immunohistochemistry and flow cytometry analysis from harvested splenocytes. We found that spleens from Mx1-Cre *Ezh2*^Y641F^ mice had proportionally lower number of B cells and a substantial increase in Ter119+ erythrocytes (**Fig. 4d**). Immunohistochemistry identified ectopic hematopoietic features consistent with extramedullary hematopoiesis (**Fig. 4c**), which is proliferation of hematopoietic tissues outside of the bone marrow. Flow cytometry analysis identified significant enrichment for hematopoietic progenitor populations in the spleens (**Fig. 4e**).

**Figure 4.**
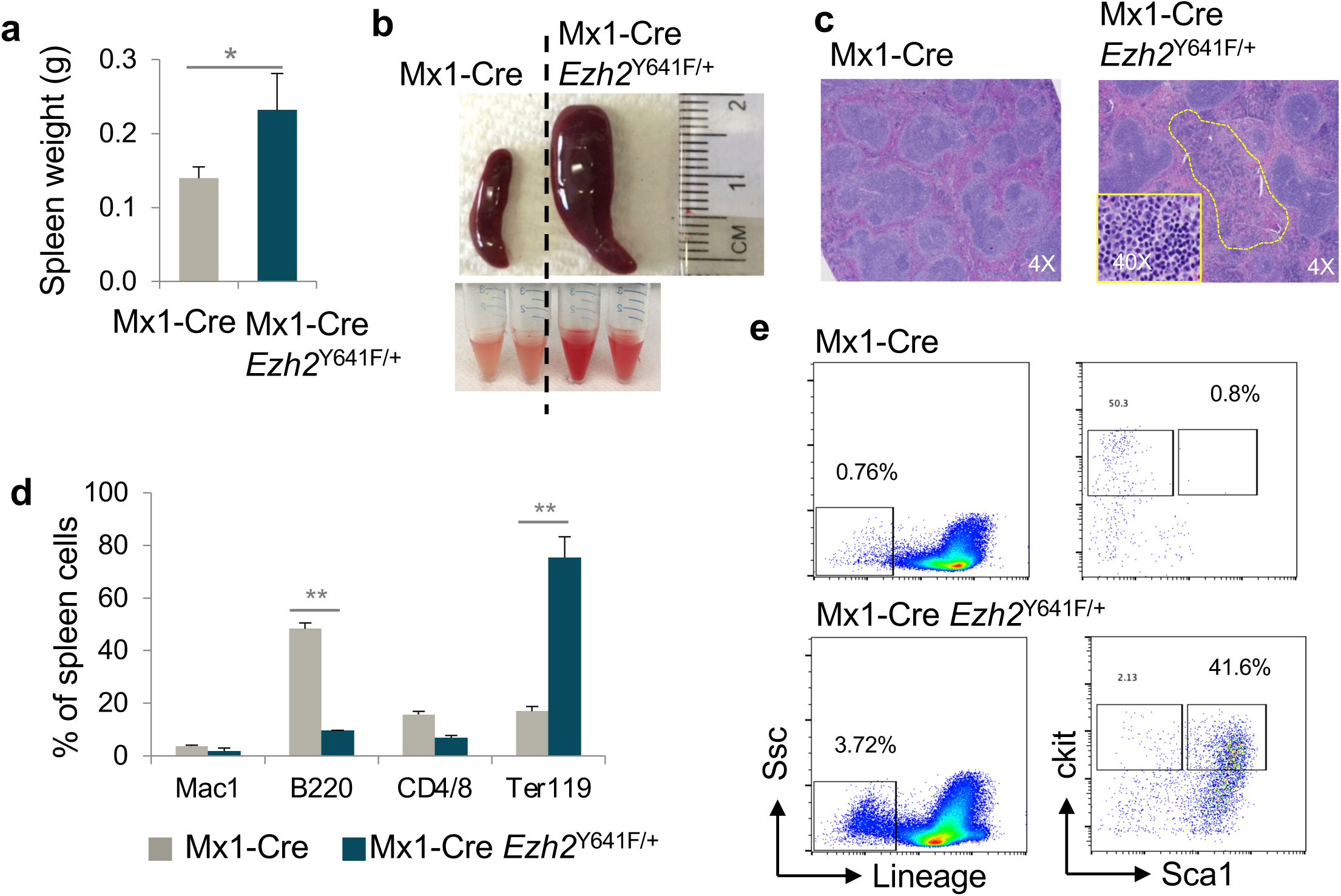
Extramedullary hematopoiesis in Mx1-Cre *Ezh2*^Y641F^ mice. **a.** Spleen weight comparison in Mx1-Cre *Ezh2*^Y641F^ versus Mx1-Cre-only control mice, n>8-10 per group. **b.** Representative images of spleen size (top) and cells in HBSS suspension (bottom). **c.** H&E-stained spleen sections showing extramedullary hematopoiesis in Mx1-Cre *Ezh2*^Y641F^ and Mx1-Cre control mice. **d.** Flow cytometry of blood lineage markers (B220, CD4, CD8, Mac1, Ter119) in spleen cells from Mx1-Cre *Ezh2*^Y641F^ and Mx1-Cre mice, n = 4-6. **e.** Flow cytometry analysis of hematopoietic progenitor populations in spleens from Mx1-Cre *Ezh2*^Y641F^ and Mx1-Cre control mice; representative of three independent experiments. Error bars represent standard deviation, * p<0.01, **p<0.001.

### Transcriptomic and pathway differences in B cell progenitors isolated from Mx1-Cre and CD19-Cre *Ezh2*^Y641F^

The findings above suggest that B cell progenitors in Mx1-Cre *Ezh2*^Y641F^ mice exhibit significant intrinsic differences compared to those from CD19-Cre mice, and that the timing of this mutation must be critical for its oncogenic activity. To better understand the differences in cell intrinsic properties of B cells isolated from the two different Cres, we performed global gene expression analysis by RNA-seq (**Fig. 5a**). Since we observed changes in B cell differentiation at progenitor stages with expression of *Ezh2*^Y641F^, we assessed differences of pro-B and pre-pro-B cells from the two different Cre crosses. To minimize experimental variation, we bred the same *Ezh2*^Y641F^ (Cre negative) male with different females, CD19-Cre or Mx1-Cre at the same time in the same cage, with the females separated before giving birth. At 12 weeks old, all mice from the Mx1-Cre groups were treated with poly(I:C) to induce Cre expression. Four weeks after poly(I:C) treatment, and at around 16-18 weeks old, an age where neither genotype exhibits any hematopoietic phenotypes, we isolated pro B and pre pro B cells from the bone marrow using FACS and processed them for RNA-sequencing. Ezh2 recombination was confirmed by PCR on spleen samples from all mice (**Supp. Fig. 2**)

**Figure 5.**
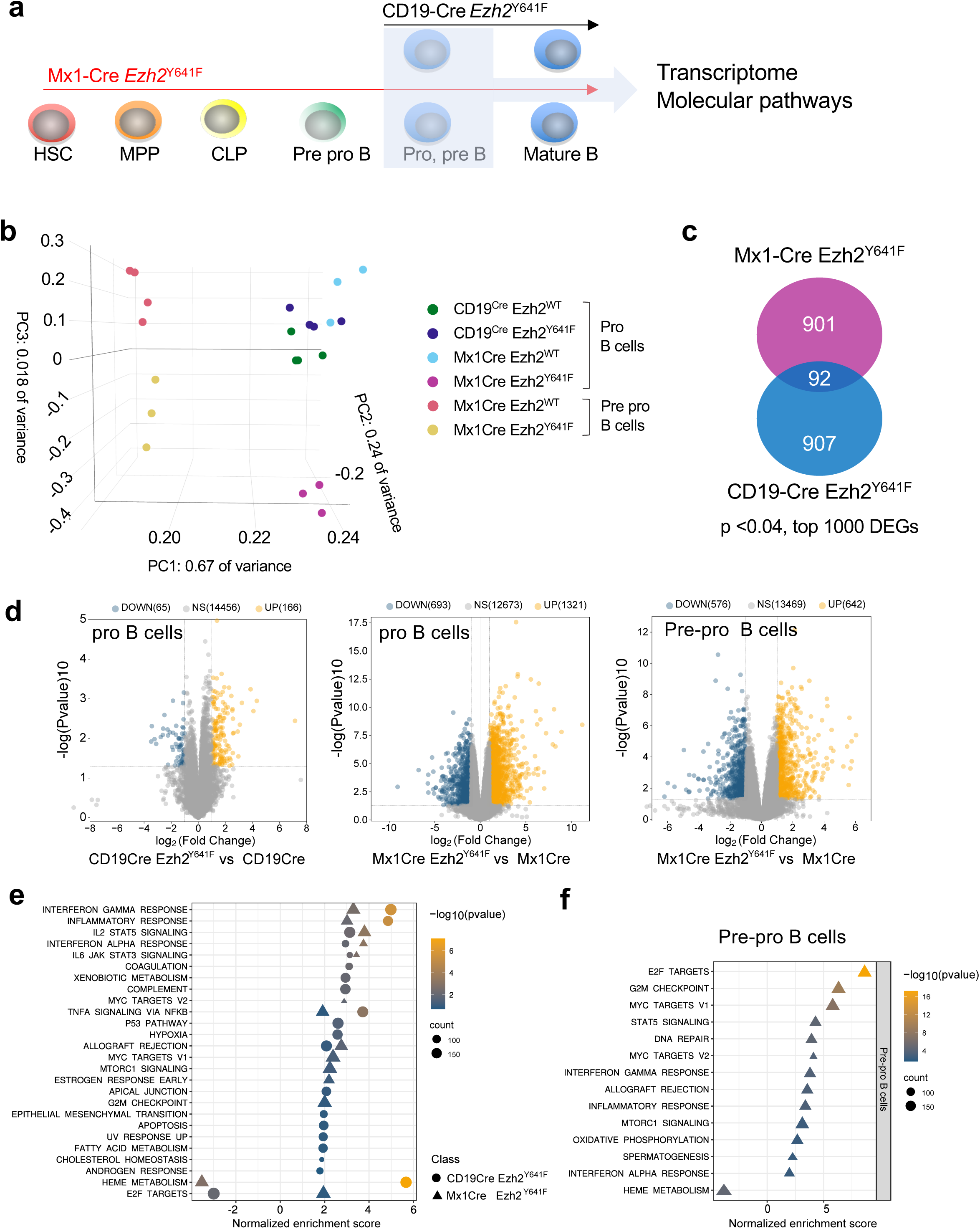
Global gene expression changes in pro and pre-pro B cells from CD19-Cre and Mx1-Cre driven expression of *Ezh2*^Y641F^. **a.** Experimental schematic: RNA-seq was performed on pre-pro and pro B cells sorted from bone marrow of *Ezh2*^WT^ and *Ezh2*^Y641F^ from Mx1-Cre and CD19-Cre models. **b.** Principal Component Analysis (PCA) plot of the RNA-seq data; pro B cells from Mx1-Cre *Ezh2*^Y641F^ and Mx1-Cre *Ezh2*^WT^ (control), n=3-4 per group. **c.** Venn diagram showing overlap of the top differentially expressed genes (DEGs) in Mx1-Cre *Ezh2*^Y641F^ vs CD19-Cre *Ezh2*^Y641F^ pro B cells. **d.** Volcano plots of DEGs in *Ezh2*^Y641F^ vs *Ezh2*^WT^ CD19-Cre pro B (left), *Ezh2*^Y641F^ vs *Ezh2*^WT^ Mx1-Cre pro B (center), and *Ezh2*^Y641F^ vs *Ezh2*^WT^ Mx1-Cre B pre pro B cells (right). **e-f.** Gene set enrichment analysis (GSEA) of hallmark signatures in CD19-Cre (circles) and Mx1-Cre (triangles) *Ezh2*^Y641F^ pro- (e) and pre pro- (f) B cells.

Analysis of gene expression data revealed both shared and distinct gene expression patterns between pro and pre-pro-B cells from the two Cre lines. Principal component analysis distinguished pro vs pre-pro B cells (PC1) and separated *Ezh2*^WT^ from *Ezh2*^Y641F^ (PC3) (**Fig. 5b)**. The separation between *Ezh2*^WT^ and *Ezh2*^Y641F^ in PC3 was more pronounced in Mx1-Cre mice compared to CD19-Cre, consistent with greater changes in gene expression with the Mx1-Cre crosses. Furthermore, the overlap in differentially expressed genes (p<0.04, top 1000 genes) was only 10% between the two crosses in pro B cells (**Fig. 5c)**. Notably, expression of *Ezh2*^Y641F^ via Mx1-Cre resulted in significantly more differentially expressed genes compared to CD19-Cre (**Fig. 5d**). Despite the limited overlap at the differentially expressed gene level, gene set enrichment analysis (GSEA) revealed significant overlap in hallmark signatures between the two Cre lines expressing *Ezh2*^Y641F^. Common signatures included enrichment for both type I and II interferon signaling, IL2-STAT5 signaling and TNF-alpha signaling (**Fig. 5e-f**). Overlapping signatures using KEGG analysis also included known EZH2 targets consistent with prior studies^36^.

We also identified distinct gene expression signatures for each Cre line. For example, BMP2 targets were enriched in the Mx1Cre *Ezh2*^Y641F^ pro B cells, while HOXA9-MEIS targets were significantly enriched in the CD19-Cre *Ezh2*^Y641F^ pro B cells (**Supp. Fig. 3**). Both pathways are directly involved in B cell differentiation and transformation and could contribute to the different phenotypes we observe with expression of mutant *Ezh2*^Y641F^ at different stages of B cell development.

Additionally, some hallmark signatures were enriched in opposite directions between the two Cre lines. These included the hallmark E2F-targets signature, which was enriched in the Mx1-Cre but depleted in the CD19-Cre (**Fig. 5e-f**), and heme metabolism, which was enriched in CD19-Cre but depleted in Mx1-Cre crosses (**Fig. 5e-f**, **Fig. 6b**). These signatures may underlie defects we observe in B cell development and differentiation, with E2F influencing cell cycle regulation, and heme metabolism via association with the Bach transcription factors ^37^ that control B cell differentiation.

**Figure 6.**
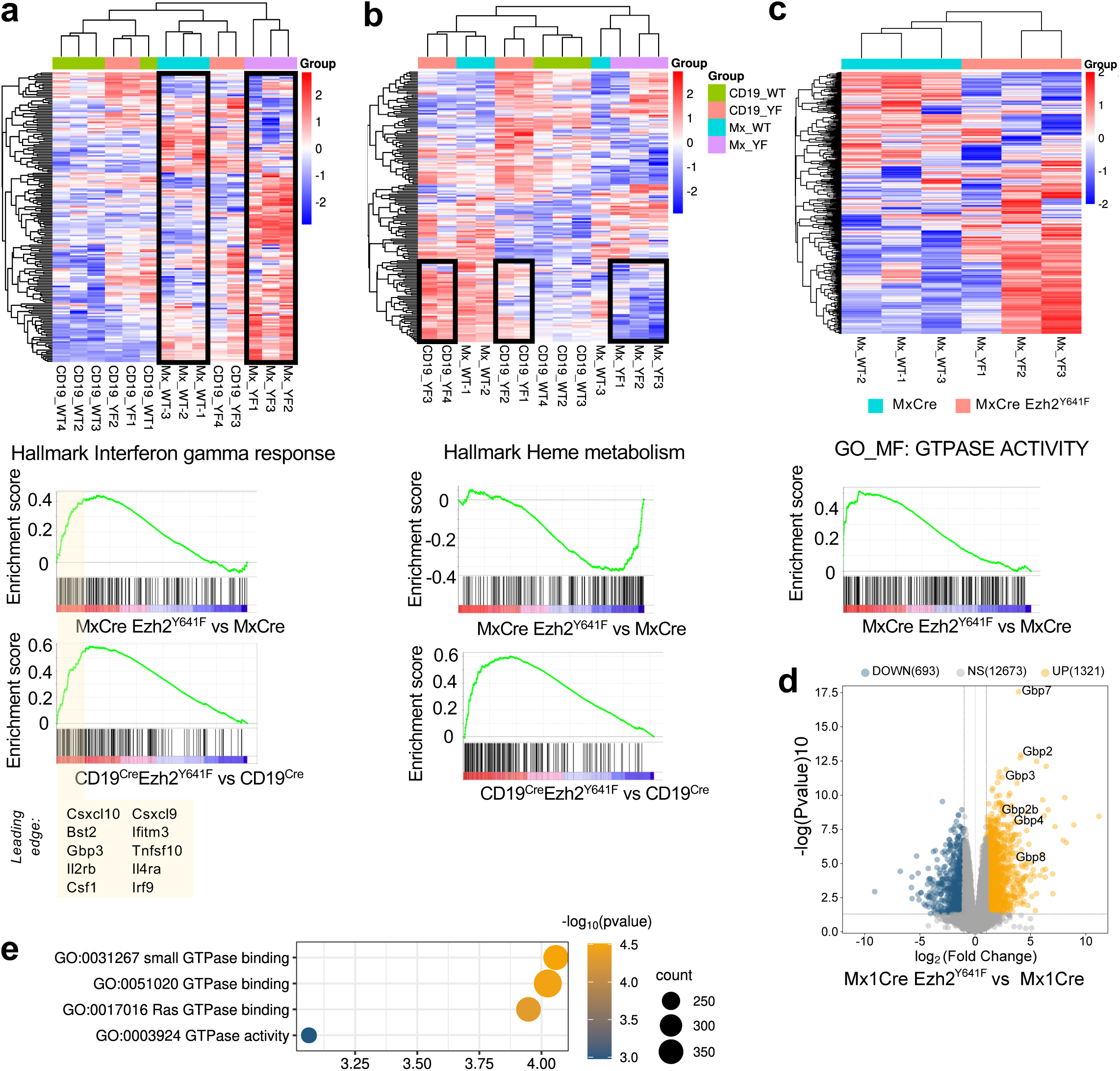
Interferon, heme metabolism and GTPase activity gene expression signatures in *Ezh2*^Y641F^ pro B cells. **a.** Top: Heatmap of interferon gamma response signature enriched in Mx1-Cre *Ezh2*^Y641F^ compared to Mx1-Cre *Ezh2*^WT^ controls; bottom: enrichment curve and genes within the leading edge. **b.** Top: Heatmap of heme metabolism signature, enriched in CD19-Cre *Ezh2*^Y641F^ and depleted in Mx1-Cre *Ezh2*^Y641F^pro B cells; bottom: enrichment curves. **c.** Top: Heatmap GTPase Activity genes, enriched in Mx1-Cre *Ezh2*^Y641F^ pro B cells compared to Mx1-Cre *Ezh2*^WT^ control; bottom: enrichment curve. **d.** Volcano plot highlighting upregulated GBP genes in Mx1-Cre *Ezh2*^Y641F^ vs Mx1-Cre *Ezh2*^WT^ control pro B cells. **e.** GSEA analysis of GTPase-related Gene Ontology (GO) terms in Mx1-Cre *Ezh2*^Y641F^ vs Mx1-Cre *Ezh2*^WT^ control pro B cells.

### Mx1-Cre *Ezh2*^Y641F^ transcriptomic changes in B cells are initiated during earlier developmental stages

To determine whether the transcriptomic changes in the pro-B cells of Mx1-Cre crosses were initiated earlier in development, we analyzed gene expression changes in the earlier, <u>pre-pro B cell stage</u>. Interestingly, the transcriptomic changes in pre-pro B cells are very similar to those in pro B cells. These include enrichment for E2F targets, MYC targets, Interferon gamma response, in addition to depletion of heme metabolism (**Fig. 5f**). This suggests that the developmental timing and cellular context shape the molecular outcomes of *Ezh2*^Y641F^ expression.

### Guanylate Binding Proteins (GBPs) are upregulated in Mx1-Cre *Ezh2*^Y641F^ pro B cells

Among the genes upregulated in Mx1-Cre *Ezh2*^Y641F^ pro B cells, a cluster of interferon-regulated GTPases known as Guanylate Binding Proteins (GBPs), stood out. These GBPs - Gbp2, 3, 4, 7 and 8 – are located within the same chromosomal region, implying potential epigenetic co-regulation (**Fig. 6c-d**). The GBP genes were also part of the leading edge of several gene expression signatures characterized by GTPase activity (**Fig. 6a, e**). GBP overexpression has mostly been associated with immune response to infections ^38,39^. In hematopoiesis, GBPs induce apoptosis in leukemia cells via interactions with MCL-1, and are associated with better prognosis^40^. In hematopoietic cancers, GBP3, 4 and 7 combined tend to be overexpressed or amplified in 14% of AML patients (**Supp. Fig. 4a**). Their expression is also correlated with expression of major IFN regulators such as STAT1 and IRF1 (**Supp. Fig. 4b-c**). In melanoma, GBP activity has been linked to anti-tumor immunity, and some GBP genes are used as prognostic biomarkers^41^.

Elevated GBP expression is common in many human cancers, with GBP1, 2, and 3 most highly expressed in cell lines from the ENCODE database (**Fig. 7a**). Expression of GBP genes appears to be modulated on EZH2 activity. For example, in human B cell lymphoma cell lines, pharmacological EZH2 inhibition resulted in significant upregulation of several GBP genes (**Fig. 7b**). Furthermore, analysis of ChIP-seq data from prior studies ^22^ showed increased H3K27me3 deposition across the entire *Gbp* locus in mouse centroblasts and centrocytes expressing mutant *Ezh2*^Y641F^ compared to controls expressing wild-type Ezh2 (**Fig. 7c**). Consistent with the mouse modeling data, EZH2 expression is negatively correlated with expression of several *GBP* genes in DLBCL patients (**Fig. 7d –** source: TCGA). In AML patients, high GBP expression, particularly GBP2, is associated with lower overall survival (TCGA) (**Fig. 7e**). These data suggest that GBP levels could serve as biomarkers or influence disease progression and outcomes in hematologic malignancies.

**Figure 7.**
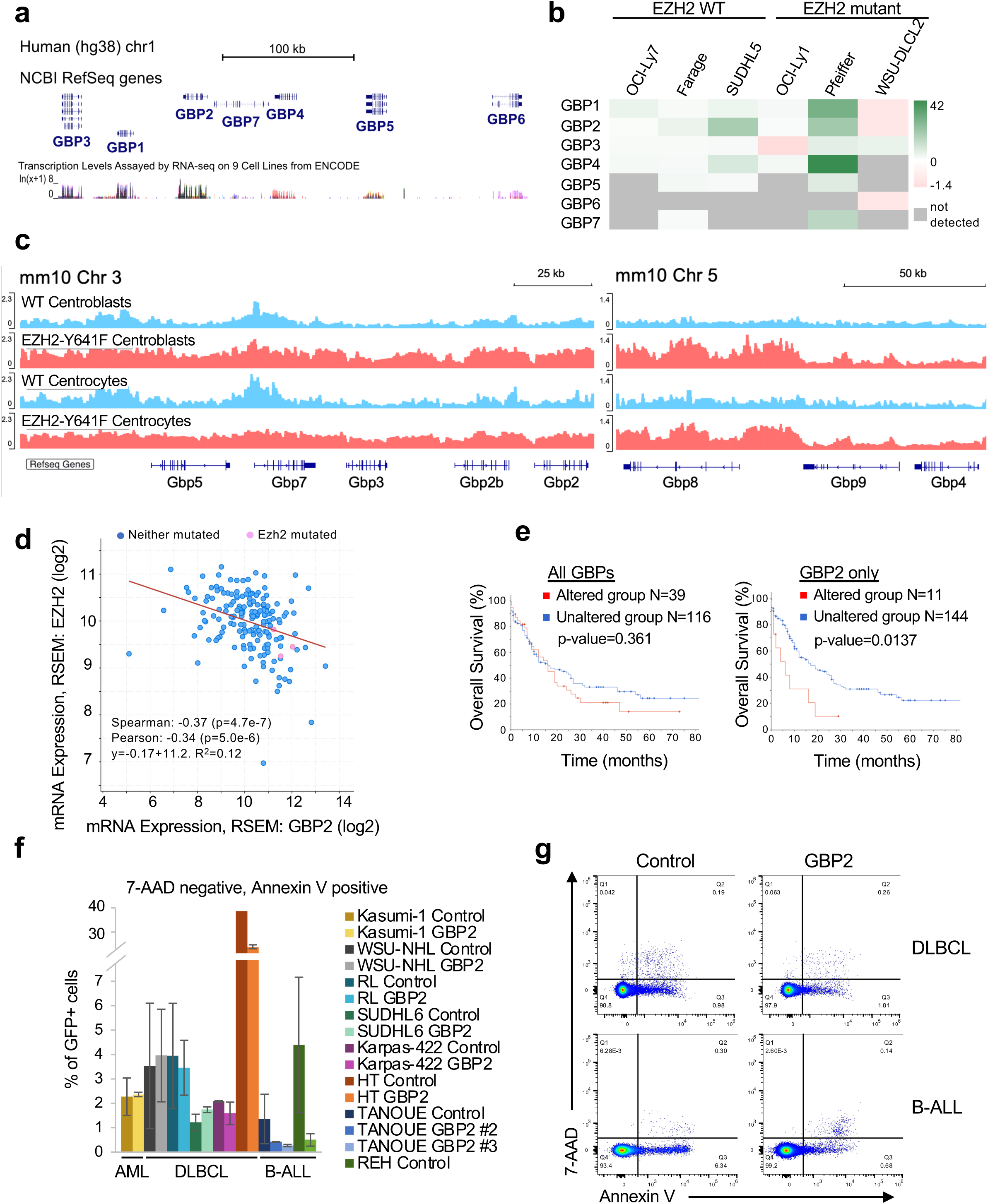
EZH2-mediated regulation of the GBP locus and its role in leukemia/lymphoma cell survival. **a.** The human GBP gene cluster and transcript levels in ENCODE RNA-seq data various cell lines (K562, GM12878, H1-hESC, HeLa-S3, HepG2, HSMM, HUVEC, NHEK, NHLF). **b.** GBP expression fold change post EZH2 inhibition with GSK343 in human lymphoma cell lines (data from GSE45982 ^8,14^). **c.** H3K27me3 ChIP-seq tracks in *Ezh2*^WT^ or *Ezh2*^Y641F^ mouse centroblasts and centrocytes at *Gbp* loci (data from GSE138036 ^22^). **d.** Correlation of *EZH2* and *GBP2* mRNA expression in AML patients (TCGA data analyzed via cBioPortal). **e.** Overall survival in AML patients with altered *GBP* expression (all GBPs vs GBP2 specifically). **f.** Flow cytometry analysis of apoptosis in GBP2 overexpressing vs and control cells (7-AAD and Annexin V staining). Results from two independent experiments. **g.** Representative flow cytometry plots from (f). Error bars represent standard deviation.

To examine whether GBP overexpression impacts survival in different types of hematopoietic malignancies, we overexpressed human *GBP2*, one of the most highly upregulated *Gbp* genes in *Ezh2*^Y641F^ pro B cells, in two B cell progenitor (BCP) leukemia cell lines, five DLBCL lines and one AML cell line, and assayed rates of apoptosis (**Supp. Fig. 4d**). While we did not observe a significance change in apoptosis in AML and DLBCL lines, we observed a reduction in apoptosis in the two pre-BCP leukemia cell lines, suggesting a context-dependent role in apoptosis mechanisms (**Fig. 7f-g**). Since *Gbp* genes appear to be co-regulated, upregulation of multiple *Gbp* genes may be required to elicit a more significant effect.

## Discussion

In this study, we provide novel insights into the complex role of *EZH2*^Y641F^ mutations across various stages of hematopoietic development and different cancer types. Using a conditional *Ezh2*^Y641F^ allele with tissue-specific Cre drivers, we demonstrated that the phenotypic and oncogenic consequences of *Ezh2*^Y641F^ are strongly influenced by developmental timing and cellular context at the time of expression. These findings extend prior studies, which primarily focused on the mutation’s role in mature B-cell lymphomas and provide insights into its broader effect during hematopoiesis and malignant transformation.

The varied outcomes with the different Cre drivers that we observe in this study underscore the importance of cell type and timing in shaping the effects of *Ezh2*^Y641F^ mutations. While CD19-Cre *Ezh2*^Y641F^ mice developed B-cell lymphoma, consistent with previous reports, Mx1-Cre, Ubc-Cre^ERT2^, and Prx1-Cre models showed significant hematopoietic defects without malignancy. These differences suggest that *Ezh2*^Y641F^ requires a permissive cellular context to drive transformation, which may include factors specific to mature B cells. The absence of malignancy in Mx1-Cre mice despite persistent *Ezh2*^Y641F^ expression in B-cell progenitors supports the idea that early disruptions to the bone marrow microenvironment may preclude the acquisition of secondary mutations necessary for transformation.

Hematopoietic defects such as bone marrow failure, extramedullary hematopoiesis, and blocks in B cell differentiation at the pre-B cell stage were seen across all Cre drivers (Ubc-, Prx1-, and Mx1-Cre models). These results suggest that there may be a unique vulnerability within hematopoietic development where *Ezh2*^Y641F^ mutations exert critical effects. This observation aligns with the higher frequency of *Ezh2*^Y641^ mutations in hematopoietic malignancies compared to solid tumors.

Bone marrow failure was a shared phenotype across most Cre crosses, particularly Mx1-Cre, Prx1-Cre, and Ubc-Cre^ERT2^, with associated splenomegaly and extramedullary hematopoiesis. These findings suggest that *Ezh2*^Y641F^ may disrupt hematopoietic homeostasis by impairing HSC self-renewal and differentiation, as was also evidenced by competitive bone marrow transplantation assays which test hematopoietic stem cell intrinsic properties. At the same time, the observed increase in trabecular bone mass and reduced marrow implicates altered bone remodeling, likely due to disrupted osteoclast function. *Ezh2*^Y641F^ expression in osteoclasts may interfere with normal bone resorption due to EZH2’s role in silencing transcription factors that negatively regulate osteoclastogenesis ^42–44^. This phenotype underscores the role of *Ezh2*^Y641F^ in shaping the microenvironment and influencing disease progression, and may have implications for understanding bone marrow failure syndromes and the interplay between hematopoietic cells and the microenvironment.

The defect in hematopoietic stem cell function is also at odds with the presence of *Ezh2*^Y641F^ mutations in AML patients. This raises questions about the role of Ezh2 and *Ezh2*^Y641F^ mutations in AML, which is thought to develop from the accumulation of genetic mutations in HSCs. *Ezh2*^Y641^ mutations in AML are very infrequent and typically of low clonality (<5%), suggesting that they are not dominant oncogenic drivers and are likely outcompeted by other more potent oncogenic clones. Since *Ezh2*^Y641F^ mutations affect stem cell fitness, there must be a unique, and uncommon context in AML where these mutations may act as oncogenic drivers, or likely emerge as secondary mutations.

Transcriptomic analysis revealed both shared and distinct molecular signatures in pro- and pre-pro B cells from Mx1-Cre and CD19-Cre *Ezh2*^Y641F^ models. Enrichment of pathways such as interferon signaling, IL2-STAT5 signaling and TNF-alpha signaling in both models suggests regulation of pathways that are independent of the timing of the mutation. However, distinct pathway enrichments, including BMP2 targets in Mx1-Cre and HOXA9-MEIS targets in CD19- Cre, highlight the fact that different timing of *Ezh2*^Y641F^ expression has different effects even at the molecular and transcriptional level. These differences may depend on the pre-existing epigenetic state of the cell at the time of *Ezh2*^Y641F^ expression, and likely underlie the divergent phenotypes observed between the models.

One of the most striking findings was the upregulation of guanylate binding proteins (GBPs) in Mx1-Cre *Ezh2*^Y641F^ pro-B cells. GBPs, interferon-regulated GTPases, are implicated in immune responses and cancer prognosis, yet their role in hematopoietic malignancies is poorly understood. High GBP expression correlates with poor outcomes in AML, and our functional analysis suggests that GBPs may influence apoptosis in a lineage-specific manner. These findings point to potential roles for GBPs as biomarkers or therapeutic targets in hematopoietic malignancies.

As with any study utilizing animal models, there are inherent limitations. The absence of malignancy in certain models may not negate the mutation’s oncogenic potential in other tissues or under different conditions. Future studies should explore additional hematopoietic-specific Cre drivers, combinations with other oncogenic mutations, and longer monitoring periods to assess the full scope of the effects of *Ezh2*^Y641F^ mutations. Investigating chromatin and histone modifications post-*Ezh2*^Y641F^ expression could provide valuable insights into the mutation’s impact on epigenetic landscapes and reveal mechanisms underlying its context-dependent behavior.

Overall, this study provides new insights into the diverse roles of *Ezh2*^Y641F^ in hematopoiesis and cancer, emphasizing the importance of developmental timing and cellular context in determining phenotypic outcomes. The identification of *GBP* upregulation as a potential regulator of apoptosis in different hematopoietic lineages introduces new opportunities for future research into the epigenetic and molecular mechanisms underlying hematopoietic malignancies. While our findings advance the understanding of EZH2’s role in hematopoietic development and disease, they also underscore the need for further investigation of the interactions between *Ezh2*^Y641F^ mutations and other oncogenic pathways. These efforts will be critical for defining therapeutic strategies and improving patient outcomes.

## Acknowledgements

We thank the Siteman Flow Cytometry Core and the McDonnell Genome Institute/Genome Technology Access Center at Washington University for flow cytometry services and genome analysis respectively, and the Department of Comparative Medicine for animal maintenance and expertise. We also thank all members of the Souroullas lab for insightful discussions and critical input on data analysis and presentation. This work was supported by the US National Cancer Institute grants K22-CA229612-01 (GPS), T32 CA113275-10 (SZ), the Cancer Research Foundation, Chicago, IL (GPS), and the Leukemia Research Foundation (GPS).

## Author contributions

SMZ, SJP, SRS, J-YL, CT and GPS performed experiments and analyzed data. J-YL performed experiments while at UNC-Chapel Hill and the work is not related to her current affiliation at Ochre Bio, Taipei, Taiwan. SMZ and GPS wrote the manuscript. GPS conceptualized and supervised the project.

## Supplemental Figure Legends

**Supplemental Figure 1. Ubc-CreERT2 and Prx1-Cre *Ezh2*^Y641F^ phenotypes.**

**a.** Immunophenotyping of lymphoma cells from CD19-Cre *Ezh2*^Y641F^ mice, characterized by B220 low, IgM high, CD19+, CD43+ and CD5+.

**b.** Abnormal hind limb posture in Prx1-Cre *Ezh2*^Y641F^ mouse.

**c.** Comparison of body weight in Ubc-Cre^ERT2^ *Ezh2*^Y641F^ vs Ubc-Cre^ERT2^ control male and female mice.

**d.** Representative spleens showing splenomegaly in the Ubc-Cre^ERT2^ *Ezh2*^Y641F^ mice.

**e.** Flow cytometry analysis of spleen cells from Ubc-Cre^ERT2^ and Prx1-Cre lines with wild-type or mutant *Ezh2*^Y641F^ alleles.

**f.** Flow cytometry analysis of blood cells from Ubc-Cre^ERT2^ with wild-type or mutant *Ezh2*^Y641F^ alleles. Lineage markers used: B220 (B cells), Mac1 (myeloid cells), CD3 (T cells) and Ter119 (erythroid cells)

**Supplemental Figure 2. Confirmation of *Ezh2*^Y641F^ recombination in B cell progenitor samples used for RNA sequencing**

PCR analysis of Ezh2 recombination at the floxed locus, confirming conversion from wild-type *Ezh2* to mutant *Ezh2*^Y641F^. PCR product at 410bp indicates the wild-type allele, while a 450bp product represents the recombined mutant allele. Positive control (Pos) is an *Ezh2*^Y641F^ melanoma cell line (Souroullas *et al.* 2016), and negative control (Neg) is a no-template control.

**Supplemental Figure 3. Unique gene expression signatures in pro B cells**

**a.** GSEA analysis shows enrichment of the Lee BMP2 Targets DN signature in Mx1-Cre *Ezh2*^Y641F^ pro B cells.

**b.** GSEA analysis shows enrichment of Hallmark MYC Targets V2 signature in Mx1-Cre *Ezh2*^Y641F^ pro B cells.

**c.** GSEA analysis shows enrichment of HOXA9 and MEIS1-regulated genes in CD19-Cre *Ezh2*^Y641F^ pro B cells.

**Supplemental Figure 4. GBP gene expression in AML patients, expression correlations and GBP2 overexpression model**

**a.** Frequency of GBP alterations in AML patients (TCGA data).

**b.** ReMap Atlas showing STAT1, STAT2, STAT3, STAT5 binding, and cis-regulatory elements at the GBP2 locus, as identified by ENCODE.

**c.** Correlation analyses of GBP1 with GBP4 (left), GBP4 with STAT1 (middle), and GBP2 with IRF1 (right) in DLBCL patients (TCGA DLBCL data, n=37, analyzed in cBioPortal).

**d.** Relative quantification of GBP2 expression by qPCR in the indicated human cell lines. Expression levels were normalized to RPL27, with fold changes calculated relative to cells transfected with an empty-vector control. Data represent the average of three replicate experiments; error bars indicate standard deviation.

