## Supplemental figures for "Developmental Stage and Cellular Context Determine Oncogenic and Molecular Outcomes of *Ezh2*^Y641F^ Mutation in Hematopoiesis"

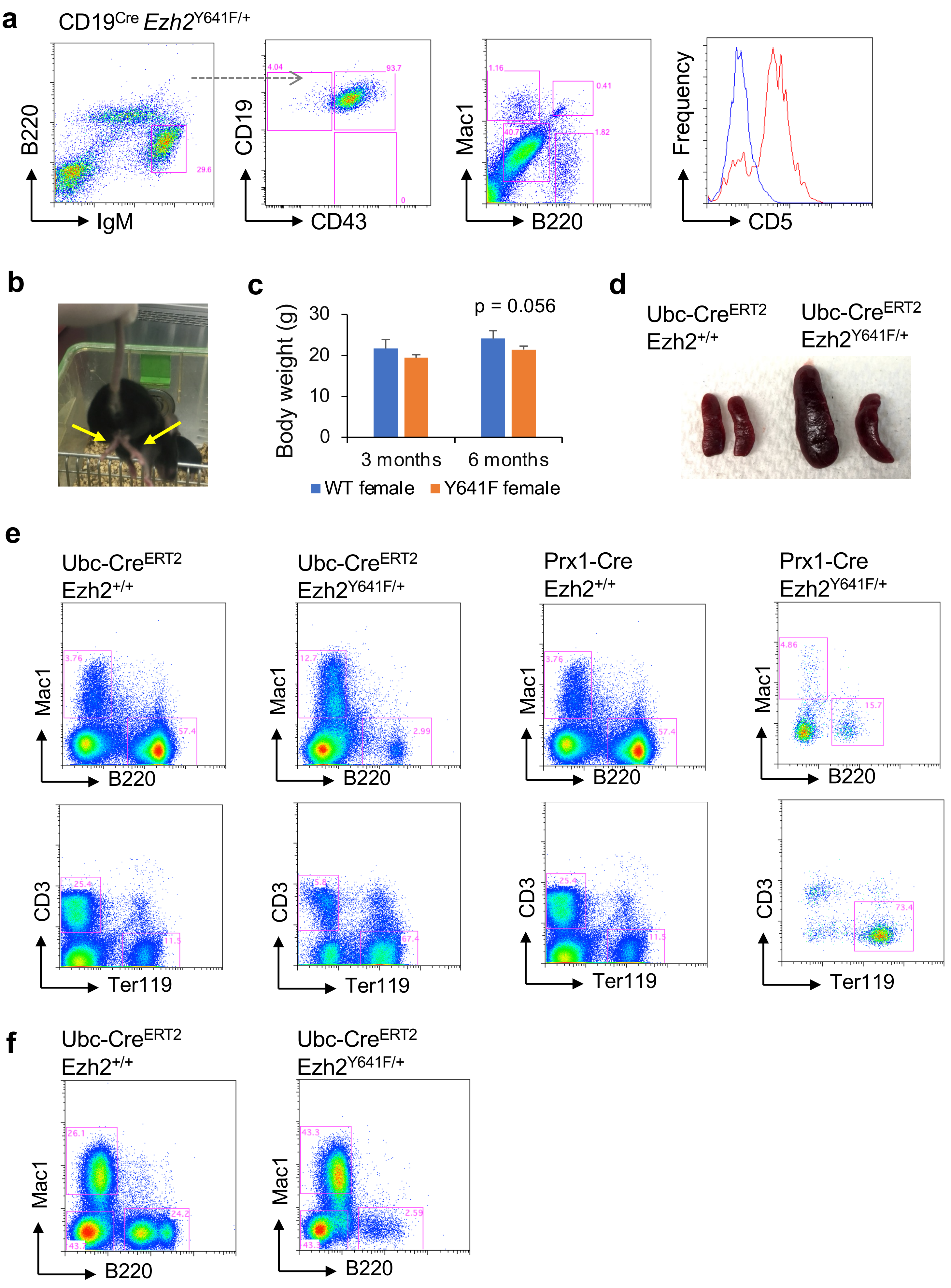


**Supp. Figure 1.** Ubc-CreERT2 and Prx1-Cre Ezh2$^{Y641F}$ phenotypes

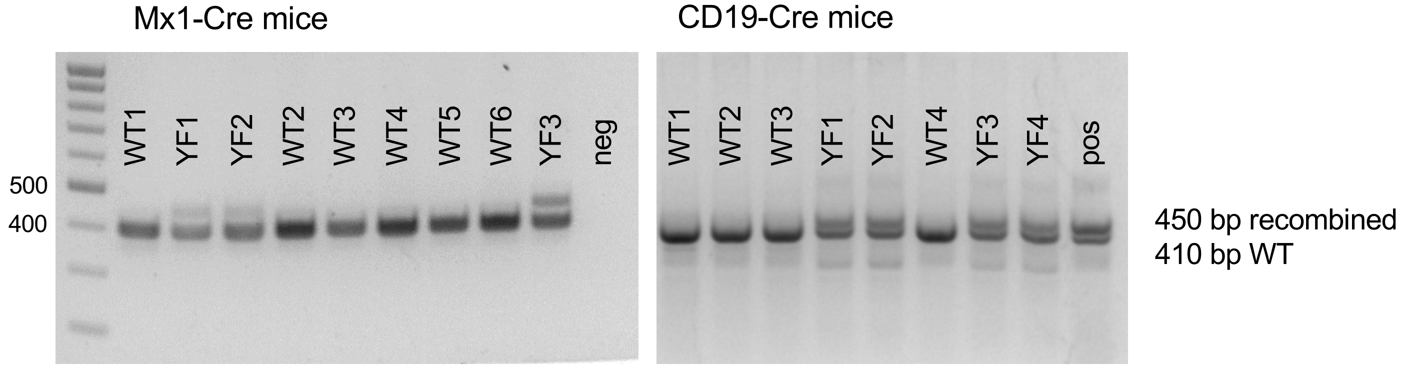


**Supp. Figure 2.** Confirmation of recombination of targeted/conditional Ezh2 allele in cells used for RNAseq

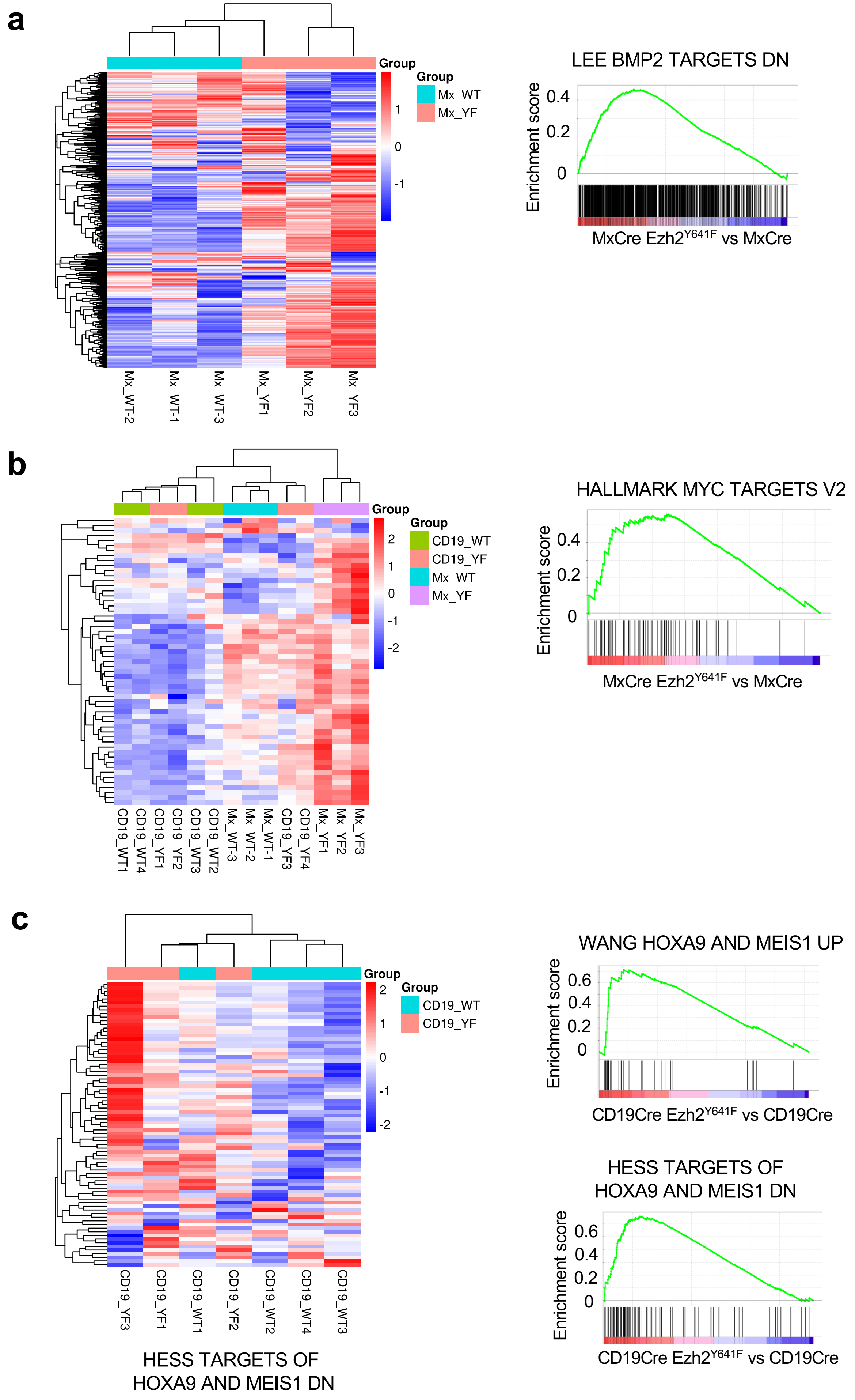


**Supp. Figure 3.** Unique gene expression signatures in pro B cells

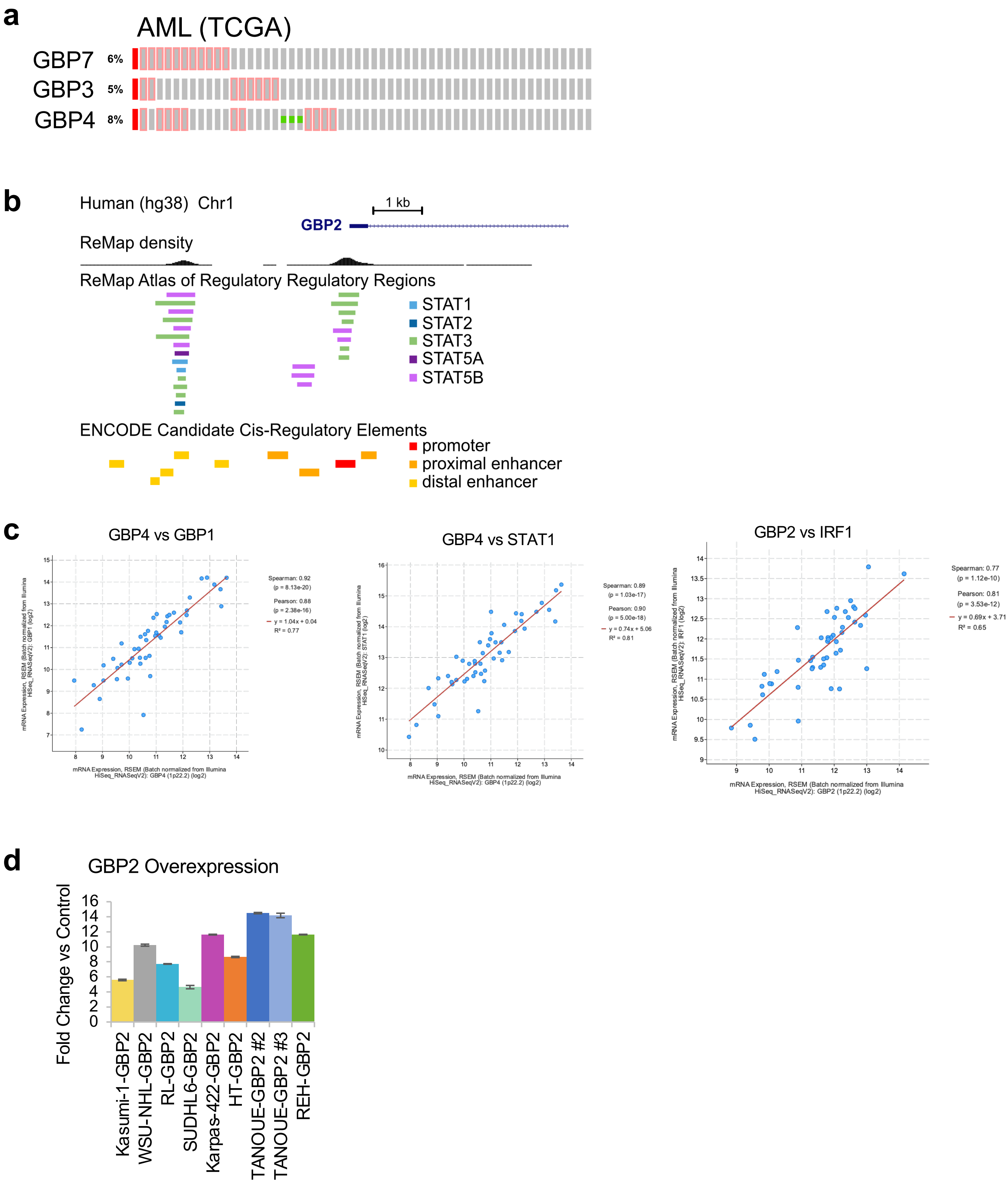


**Supp. Figure 4.** GBP gene expression in AML patients, expression correlations and GBP2 overexpression model
