## Supplementary material for "Developmental Stage and Cellular Context Determine Oncogenic and Molecular Outcomes of *Ezh2*^Y641F^ Mutation in Hematopoiesis": Supp. table 1

| **Antibodies** | **Clone** | **Source** |
| --- | --- | --- |
| CD45R/B220 | RA3-6B2 | eBioscience |
| CD3 | 145-2c11 | eBioscience |
| CD45.2 | 104 | eBioscience |
| CD4 | GK1.5 | eBioscience |
| CD8 | 53-6.7 | eBioscience |
| CD19 | eBio1D3 | eBioscience |
| CD5 | 53-7.3 | eBioscience |
| CD21 | eBio4E3 | eBioscience |
| CD43 | eBioR2/60 | eBioscience |
| IgD | 11-26 | eBioscience |
| Gr1 | RB6-8C5 | eBioscience |
| IgM | 11f41 | eBioscience |
| GL-7 | GL-7 | eBioscience |
| Mac1 | M1/70 | eBioscience |
| ckit | 2B8 | Biolegend |
| Sca1 | D7 | Biolegend |
| IL7Ra | A019D5 | Biolegend |
| Annexin V | 640920 | Biolegend |
| 7-AAD | 420403 | Biolegend |

**Supplemental Table 1.** List of antibodies used for flow cytometry.
