## Supplementary material for "Developmental Stage and Cellular Context Determine Oncogenic and Molecular Outcomes of *Ezh2*^Y641F^ Mutation in Hematopoiesis": Supp. table 2

| | |
|---|---|
| **Cre** | |
| MB182F | 5’-CAATGGTAGGCTCACTCTGGGAGATGATA-3' |
| MB182R | 5’-AACACACACTGGCAGGACTGGCTAGG-3' |
| MB183F | 5’-CAGGGTGTTATAAGCAATCCC-3' |
| MB183R | 5’- CCTGGAAAATGCTTCTGTCCG-3' |
| **Ezh2 Y641F allele** | |
| Forward | 5’-CTCTTGGGGGTCAGTCCAC-3' |
| Reverse | 5’-AAGTTGCTGAGTGTCTGGAAAA-3' |
| **Ezh2 allele recombination** | |
| Forward | 5’- ACAGAAGTGAGAGAAGCTGATC-3' |
| Reverse | 5’- GCTGCTACTATAAACAAGTC-3' |
| **Human GBP2 gene expression (Sybr green)** | |
| Forward | 5’-CAGTTGGAAGCAAGGCGAGAT-3' |
| Reverse | 5’-GCACCTCTTTGGCCTGTATCC-3' |
| **Human RPL27 gene expression (Sybr green)** | |
| Forward | 5’-ATCGCCAAGAGATCAAAGATAA-3' |
| Reverse | 5’-TCTGAAGACATCCTTATTGACG-3' |

**Supplemental Table 2.** Primer sequences
